# RNA-spray-mediated silencing of *Fusarium graminearum AGO* and *DCL* genes improve barley disease resistance

**DOI:** 10.1101/821868

**Authors:** B Werner, FY Gaffar, J Schuemann, D Biedenkopf, A Koch

## Abstract

Over the last decade, several studies have revealed the enormous potential of RNA-silencing strategies as a potential alternative to conventional pesticides for plant protection. We have previously shown that targeted gene silencing mediated by an *in planta* expression of non-coding inhibitory double-stranded RNAs (dsRNAs) can protect host plants against various diseases with unprecedented efficiency. In addition to the generation of RNA-silencing (RNAi) signals *in planta*, plants can be protected from pathogens and pests by spray-applied RNA-based biopesticides. Despite the striking efficiency of RNA-silencing-based technologies holds for agriculture, the molecular mechanisms underlying spray-induced gene silencing (SIGS) strategies are virtually unresolved, a requirement for successful future application in the field. Based on our previous work, we predict that the molecular mechanism of SIGS is controlled by the fungal-silencing machinery. In this study, we used SIGS to compare the silencing efficiencies of computationally-designed versus manually-designed dsRNA constructs targeting *ARGONAUTE* and *DICER* genes of *Fusarium graminearum* (*Fg*). We found that targeting key components of the fungal RNAi machinery via SIGS could protect barley leaves from *Fg* infection and that the manual design of dsRNAs resulted in higher gene-silencing efficiencies than the tool-based design. Moreover, our results indicate the possibility of cross-kingdom RNA silencing in the *Fg*-barley interaction, a phenomenon in which sRNAs operate as effector molecules to induce gene silencing between species from different kingdoms, such as a plant host and their interacting pathogens.

## Introduction

Diseases of cereal crops, such as Fusarium head blight caused by phytopathogenic fungi of the genus *Fusarium* and primarily by the ascomycete *Fusarium graminearum* (*Fg*), exert great economic and agronomic impacts on global grain production and the grain industry (Goswami and Kistler 2004; Kazan et al. 2012; McMullen et al. 2012). In addition to significant yield losses, food quality is adversely affected by grain contamination with mycotoxins, representing a serious threat to human and animal health (Ismaiel and Papenbrock 2015). Plant-protection and toxin-reduction strategies are presently mediated by chemical treatments. Currently, the application of systemic fungicides, such as sterol demethylation inhibitors (DMIs), is essential for controlling Fusarium diseases and to assist in reaching the maximum attainable production level of high-yield cultivars. DMI fungicides act as ergosterol biosynthesis inhibitors due to cytochrome P450 lanosterol C-14α-demethylase (CYP51) binding, which subsequently disturbs fungal membrane integrity (Kuck et al. 2012). Because of a shortage of alternative chemicals, DMIs have been used extensively in the field since their discovery in the 1970s. Therefore, it is hardly surprising that reduced sensitivity, or even resistance to DMI fungicides, has begun to develop in many plant pathogenic fungi (Yin et al. 2009; Spolti et al. 2014). These developments demonstrate an urgent need for novel strategies in pathogen and pest control.

RNAi, known as a conserved and integral part of the gene regulation processes present in all eukaryotes, is mediated by small RNAs (sRNAs) that direct gene silencing at the post-transcriptional level. Post-transcriptional gene silencing starts with the initial processing or cleavage of a precursor double-stranded (ds)RNA into short 21-24 nucleotide (nt) small-interfering RNA (siRNA) duplexes by an RNaseIII-like enzyme called Dicer (Baulcombe, 2004; Ketting, 2011). Double-stranded siRNAs are incorporated into an RNA-induced silencing complex that initially unwinds the siRNA, thereby generating an antisense (or guide) strand which base-pairs with complementary mRNA target sequences. Subsequent degradation of the targeted mRNA mediated by an RNase protein called Argonaute (AGO) prevents translation of the target transcript (Vaucheret et al., 2004; Borges and Martienssen 2015), ideally resulting in a loss of function phenotype. Therefore, RNAi has emerged as a powerful genetic tool not only in fundamental research for the assessment of gene function but also in various fields of applied research, such as agriculture. In plants, RNAi strategies have the potential to protect host plants against predation or infection by pathogens and pests mediated by lethal RNAi signals generated *in planta*, a strategy known as ‘host-induced gene silencing’ (HIGS; Nowara et al. 2010) (for review, see Koch and Kogel 2014; Yin and Hulbert 2015; Zhang et al., 2017; Qi et al., 2019; Gaffar and Koch 2019). In addition to the generation of RNA-silencing signals *in planta*, plants can be protected from pathogens and pests by spray-applied RNA biopesticides, known as spray-induced gene silencing (SIGS) (Koch et al., 2016; Wang et al., 2016; Konakalla et al., 2016; Mitter et al., 2017; Kaldis et al., 2018; Koch et al. 2019). Regardless of how target-specific inhibitory RNAs are applied (i.e. endogenously or exogenously), the use of HIGS and SIGS technologies to control *Fusarium* species have been shown to be a potential alternative to conventional pesticides (Koch et al. 2013; Ghag et al. 2014; Cheng et al. 2015; Hu et al. 2015; Chen et al. 2016; Pareek and Rajam 2017; Bharti et al. 2017; Baldwin et al. 2018; Koch et al. 2018; Koch et al. 2019), supporting the notion that RNAi strategies may improve food safety by controlling the growth of phytopathogenic, mycotoxin-producing fungi (reviewed by Machado et al. 2017; Majumdar et al. 2017).

Despite the notable efficiency RNAi-based technology holds for agriculture, the mechanisms underlying HIGS and SIGS technologies are inadequately understood. There is little information regarding the contribution of either plant- or fungal-silencing machinery in cross-species RNA silencing (i.e. plant and fungus) or how inhibitory RNAs translocate from the plant to the fungus after its transgenic expression or spray application. Whereas HIGS is virtually based on the plant’s ability to produce mobile siRNAs (through plant Dicers [DCLs]), the mechanism of gene silencing by exogenously delivered dsRNA requires the fungal RNAi machinery, mainly fungal DCLs (Koch et al. 2016, Gaffar et al. 2019). Interestingly, recent studies revealed that AGO and DCL proteins of *Fg* contribute to vegetative and generative growth, disease development, mycotoxin production, antiviral response and sensitivity to environmental RNAi (Kim et al. 2015; Son et al. 2017; Yu et al. 2018; Gaffar et al. 2019). In *Fg*, two Dicer proteins (*Fg*DCL1 and *Fg*DCL2) and two AGO proteins (*Fg*AGO1 and *Fg*AGO2) were identified (Chen et al. 2015). Characterization of those RNAi core components revealed functional diversification, as *Fg*AGO1 and *Fg*DCL2 were shown to play important roles in hairpin-RNA-induced gene silencing (Chen et al. 2015). In addition, we recently demonstrated that *Fg*AGO2 and *Fg*DCL1 are required for sex-specific RNAi (Gaffar et al. 2019). Moreover, *Fg*AGO2 and *Fg*DCL1 participate in the biogenesis of perithecium-specific microRNAs (Zeng et al. 2018).

Notably, we previously demonstrated that *Fg*DCL1 is required for SIGS-mediated *Fg* disease resistance (Koch et al. 2016). However, further analysis of *Fg* RNAi KO mutants revealed that all tested mutants were slightly or strongly compromised in SIGS, whereas *FgCYP51* target gene expression was completely abolished *in Δdcl2* and *Δqip1* mutants (Gaffar et al. 2019). Together, these studies indicate a central role of RNAi pathways in regulating *Fg* development, pathogenicity and immunity. Consistent with this notion, we assume that *Fg* RNAi components represent suitable targets for RNA spray-mediated disease control. To determine this, we generated different dsRNA constructs targeting *FgAGO* and *FgDCL* genes that were sprayed onto barley leaves. We also compared two different dsRNA design strategies; thus, we used a tool-based prediction of suitable dsRNA construct sequences versus a manual construct design related to current design principles and experiences. Notably, tool- and manually-designed RNAi constructs differ also in the length of the dsRNA molecules that were applied by the topical spray of barley leaves.

## Material and Methods

### Construction of AGO1, AGO2, DCL1 and DCL2 templates and synthesis of dsRNA

Primers were designed to generate PCR amplicons 658–912 bp in length for the manually-designed construct or 173–193 bp in length for the tool-designed construct (Zhao Bioinformatics Laboratory tool; http://plantgrn.noble.org/pssRNAit/), corresponding to exons of selected target genes, in which *Fg* represents *Fusarium graminearum*: *FgAGO1* (FGSG_08752), *FgAGO2* (FGSG_00348), *FgDCL1* (FGSG_09025) and *FgDCL2* (FGSG_04408) (Figure S1-4). The target gene sequences were amplified from *Fg* wt strain IFA65 cDNA using target-specific primers (Table S1). The length of manually selected sequences were 658 bp from *FgAGO1*, 871 bp from *FgAGO2*, 912 bp from *FgDCL1* and 870 bp from *FgDCL2*, while the respective tool selected sequences were 173 bp, 192 bp, 182 bp and 193 bp.

The construction of pGEMT plasmids comprised of the tool- and manually-designed target sequences was performed using restriction enzyme-cloning strategies. The first step in constructing pGEMT plasmids containing manually-designed double targets was to amplify target sequences of *AGO1, AGO2, DCL1* and *DCL2* from the confirmed plasmids with primers containing restriction sites (Table S1). The manually designed dsRNA targeting *FgAGO1* and *FgAGO2* had a length of 1529 bp and was therefore named ago1/ago2_1529nt. According to this scheme the other manually-designed dsRNAs were named ago1/dcl1_1570nt, ago1/dcl2_1528nt, ago2/dcl1_1783nt, ago2/dcl2_1741nt and dcl1/dcl2_1782nt. Briefly, an *AGO2* PCR fragment was inserted between NotI and NdeI restriction sites of pGEMT plasmids containing *AGO1* or *DCL1* target sequences to generate ago1/ago2_1529nt and ago1/dcl2_1528nt constructs. The PCR fragment of *AGO1* was inserted between NotI and NdeI restriction sites of pGEMT plasmids containing the *DCL1* target sequence to construct ago1/dcl1_1570nt target plasmid. The other manually designed constructs (ago1/dcl2_1528nt, ago2/dcl2_1741nt and dcl1/dcl2_1782nt) were generated following the same procedure as described above: DCL2 PCR fragments were inserted in the AGO1 background (using NotI and NdeI), in AGO2 (using NotI and BstXI) and in DCL1 (using NotI and SalI). To construct pGEMT plasmids containing tool-designed target sequences (ago1/ago2_365nt, ago1/dcl1_355nt, ago2/dcl1_374nt, ago1/dcl2_366nt), the single targets were amplified using primers containing a restriction site (Table S1), as described above. A tool-designed sequence of *DCL1* was inserted between NotI and SalI restriction sites of the pGEMT plasmid containing *AGO1* and *AGO2* targets to generate ago1/dcl1_355nt and ago2/dcl1_374nt constructs, respectively. The *DCL2* fragment was inserted between the NotI and SalI restriction sites of the pGEMT plasmid containing the *AGO1* sequence to construct ago1/dcl2_366nt. Finally, *AGO2* was inserted between the NotI and SalI restriction sites of the pGEMT plasmid containing the *AGO1* target sequence to generate an ago1/ago2_365nt construct.

MEGAscript Kit High Yield Transcription Kit (Ambion) was used for dsRNA synthesis by following the manufacturers’ instructions using primers containing a T7 promoter sequence at the 5′ end of both forward and reverse primers (Table S1).

### Spray application of dsRNA on barley leaves

The second leaves of 2- to 3-wk-old barley cultivar Golden Promise were detached and transferred to square Petri plates containing 1% water-agar. The dsRNA was diluted in 500 μl of water to a final concentration of 20 ng μl^-1^. For the Tris-EDTA (TE) control, TE buffer was diluted in 500 μl of water, corresponding to the amount used for dilution of the dsRNA. The typical RNA concentration after elution was 500 ng μl^-1^, representing a buffer concentration of 400 μM of Tris-HCL and 40 μM of EDTA in the final dilution. Leaves were sprayed using a spray flask as described earlier (Koch et al., 2016). Each plate containing ten detached leaves was evenly sprayed (3-4 puffs) with the different tool- and manually-designed dsRNAs or TE buffer and subsequently kept at room temperature. Forty-eight hours after spraying, leaves were drop-inoculated with three 20-µl drops of *Fg* suspension containing 5 × 10^4^ conidia ml^-1^ water. After inoculation, plates were closed and incubated for five days at room temperature. The relative infection of the leaves was recorded as the infection area (by determining the size of the chlorotic lesions) relative to the total leaf area using ImageJ (Schneider et al., 2012) software. We produced four biological replicates for independent sample collection.

### Fungal transcript analysis

To assess the silencing of the *FgAGO* and *FgDCL* genes, mRNA expression analysis was performed using quantitative reverse-transcription PCR (qRT-PCR). RNA extraction from the diseased leaves was performed with TRIzol (Invitrogen) following the manufacturer’s instructions. Freshly extracted mRNA was used for cDNA synthesis using a qScript™ cDNA kit (Quantabio). For qRT-PCR, 10 ng of cDNA was used as a template with the reactions run in a QuantStudio 5 Real-Time PCR system (Applied Biosystems). Amplifications were performed in 7.5 μl of SYBR^®^ Green JumpStart Taq ReadyMix (Sigma-Aldrich) with 5 pmol of oligonucleotides. Each sample underwent three technical repetitions. Primers were used for studying expressions of *FgAGO* and *FgDCL* genes with reference to the *Elongation factor 1-alpha* (*EF1-a*) gene (FGSG_08811) and *ß-tubulin* (Table S1). After an initial activation step at 95°C for 5 min, 40 cycles (95°C for 30 sec, 57°C for 30 sec, 72°C for 30 sec) were performed. Cycle threshold values were determined using the 7500 Fast software supplied with the instrument. Levels of *FgAGO* and *FgDCL* transcripts were determined via the 2^-ΔΔCt^ method (Livak and Schmittgen 2001) by normalizing the amount of target transcript to the amount of the reference transcripts of the *EF1-a* and *ß-tubulin*.

### siRNA prediction

Sequences of the single manually- and tool-designed dsRNA constructs for each gene, *FgAGO1, FgAGO2, FgDCL1* and *FgDCL2*, were split into k-mers of 21 bases and mapped to the coding sequences of the 4 *FgAGO* and *FgDCL* genes. The efficient siRNAs were calculated on the basis of the thermodynamic properties of the siRNA-duplex, the 5’-nucleotide of the guide strand and the target site accessibility based on the default parameters of the SI-FI software tool (http://labtools.ipk-gatersleben.de/). These parameters were no mismatches to the target sequence, a 5’-A or -U on the potential guide strand, a higher minimum free energy (MFE) on the 5’-end of the guide strand compared to the passenger strand and good target site accessibility. To this end, the default parameters were used.

## Results

### Spray-induced gene silencing by AGO- and DCL-dsRNAs reduces *Fg* infection

We assessed whether *FgAGO* and *FgDCL* genes are suitable targets for SIGS-mediated plant protection strategies. Detached barley leaves were sprayed with 20 ng µl^-1^ dsRNA and drop-inoculated 48 h later with a suspension of *Fg* conidia. After 5 dpi, necrotic lesions were visible at the inoculation sites of leaves sprayed with TE buffer or GFP-dsRNA as negative controls. All dsRNAs reduced the infection symptoms, as revealed by significantly smaller lesions (Figure 1). Infected areas were reduced by all constructs, on average, by 50% compared to the control (Figure 1). The highest infection reduction of 60% was reached with dsRNAs targeting ago1/ago2_365nt and ago1/dcl1_1570nt (Figure 1). The lowest disease resistance efficiencies of 31% were shown for the ago2/dcl1_1783nt dsRNA construct (Figure 1).

**Figure 1:**
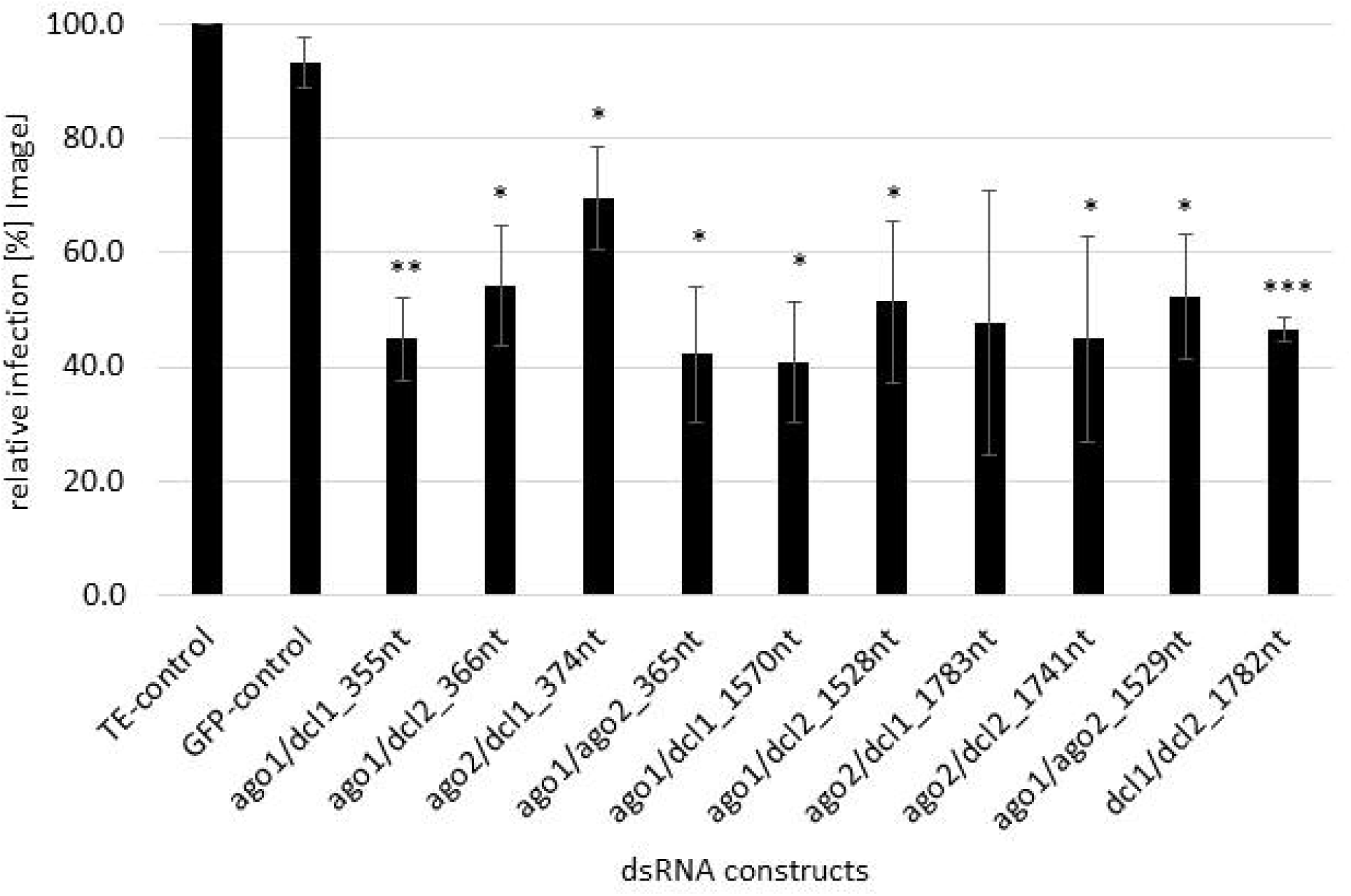
Quantification of infection symptoms of *Fg* on barley leaves sprayed with AGO/DCL-targeting dsRNAs. Detached leaves of 3-wk-old barley plants were sprayed with AGO/DCL-targeting dsRNAs or TE buffer. After 48 h, leaves were drop inoculated with 5 x 10^4^ ml^-1^ of macroconidia and evaluated for infection symptoms at 5 dpi. Infection area, shown as the percent of the total leaf area for 10 leaves for each dsRNA and the TE control relative to the infected leaf area. Bars represent mean values ± SDs of three independent experiments. Asterisks indicate statistical significance (**p<0,01; ***p< 0,001; students t-test).

### DCL-dsRNAs exhibited higher target gene silencing than AGO-dsRNAs

To analyse whether the observed resistance phenotypes were provoked by target gene silencing, we measured the transcript levels of *FgAGO* and *FgDCL* genes of *Fg* grown in the infected leaf tissue by qRT-PCR. As anticipated, the relative transcript levels of targeted genes *FgAGO1, FgAGO2, FgDCL1* and *FgDCL2* were reduced after the inoculation of leaves sprayed with the respective dsRNA constructs (Figure 2A/B), and except for *FgAGO1*, if targeted with tool-designed constructs ago1/dcl1_355nt, ago1/dcl2_366nt and ago1/ago2_365nt (Figure 2A). However, regarding those three constructs, we detected silencing effects for the second target gene, as the *FgDCL1* expression was reduced by 47%, *FgDCL2* by 44% and *FgAGO2* by 52% (Figure 2A). The most efficient construct in terms of overall target gene silencing was ago2/dcl1_374nt, which reduced the transcripts of *FgAGO2* and *FgDCL1* by 40% and 74%, respectively, compared to the TE control (Figure 2A/B).

**Figure 2:**
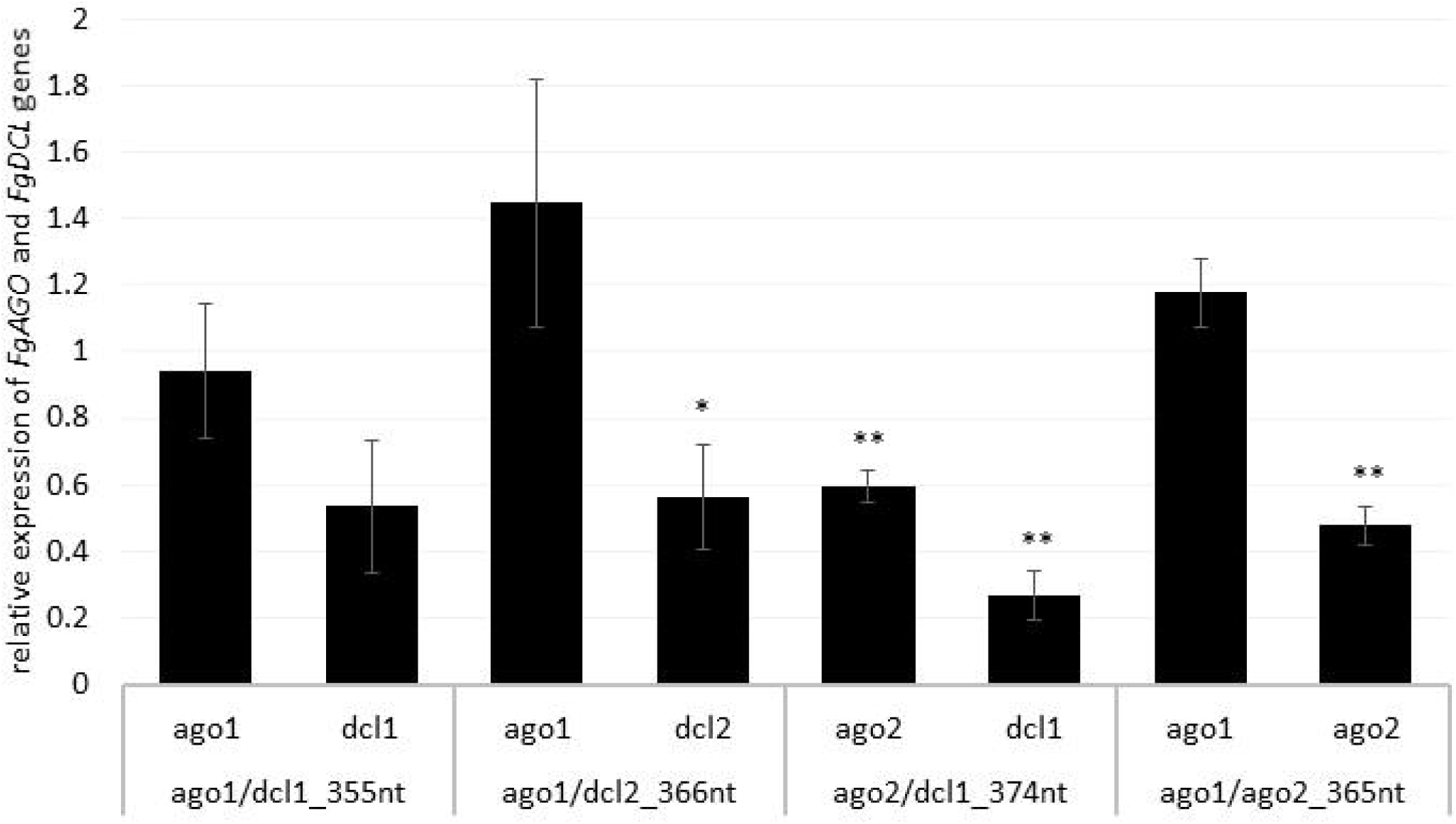
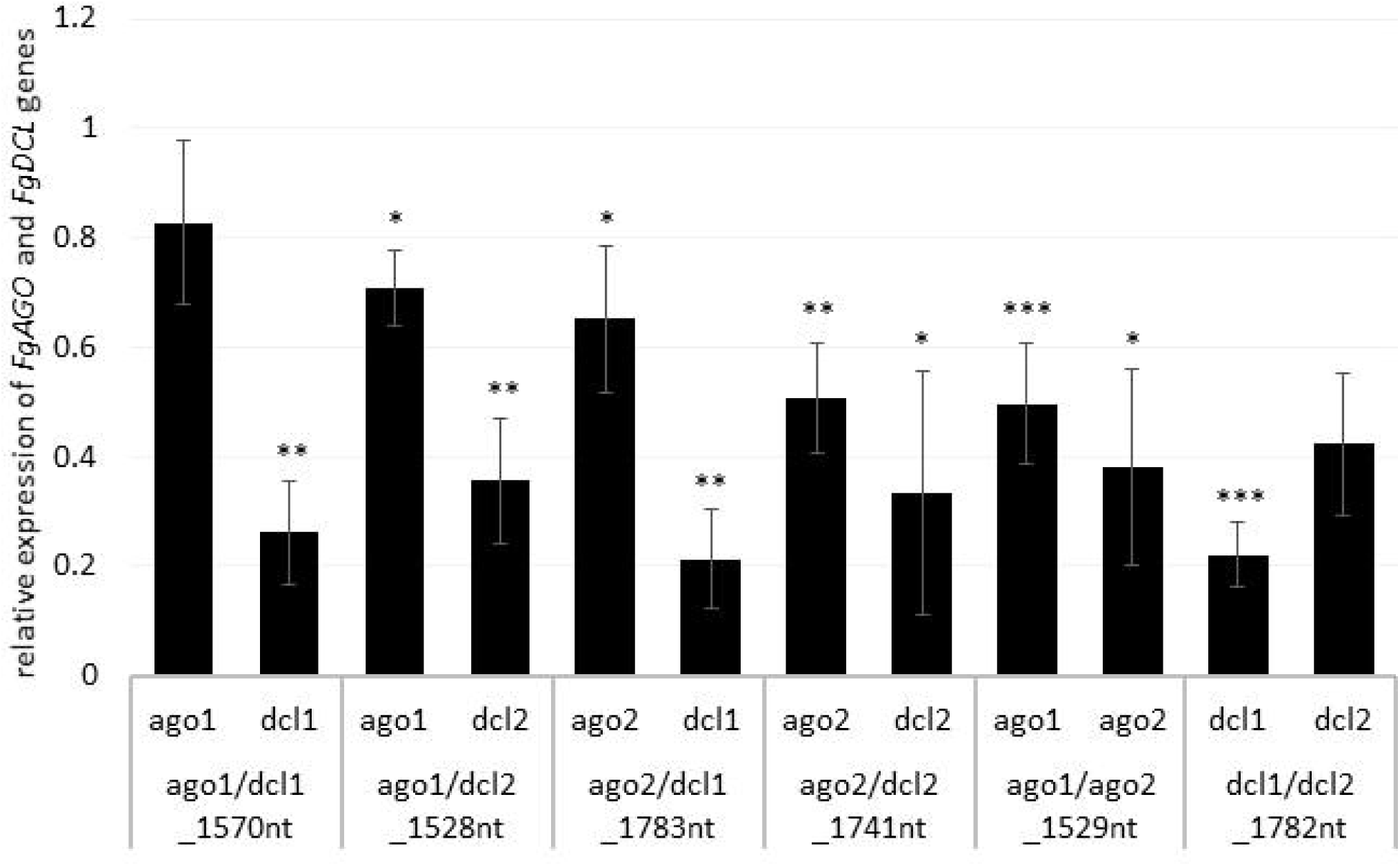
Relative expression of the respective fungal *DCLs* and *AGOs* 5 dpi on (A) tool- and (B) manually-designed-dsRNA-sprayed leaves. The expression was measured via the ΔΔ-ct method in which the expression of the respective *AGOs* and *DCLs* was normalized against the fungal reference gene *EF1α (translation elongation-factor 1 α)* and *ß-tubulin*, and this Δ-ct value was then normalized against the Δ-ct of the GFP control. Error bars represent the SE of the four independent experiments, each using 10 leaves of 10 different plants for each transgenic line. Asterisks indicate statistical significance (*p<0,05; **p<0,01; ***p < 0,001; students t-test).

Notably, if we compared the results for the tool-designed dsRNA constructs with the manually-designed dsRNAs we observed similar results for the *FgAGO1* target-silencing (Figure 2A/B). The constructs ago1/dcl1_1570nt and ago1/dcl2_1528nt reduced *FgAGO1* transcripts by only 17% and 29%, respectively (Figure 2B). Analysing the transcript levels of *FgAGO2* revealed that (1) the silencing efficiencies of ago2/dcl1_1783nt and ago2/dcl2_1741nt were higher than *FgAGO1* target silencing and (2) targeting both *FgAGO* genes with the ago1/ago2_1529nt construct resulted in 50% reduction for *FgAGO1* and 62% for *FgAGO*2. This, therefore, showed the highest overall *FgAGOs* gene silencing (Figure 2B).

Interestingly and consistent with the tool-designed target gene silencing results, we detected the strongest reduction of >70% for *FgDCL1* (Figure 2B). For example, ago2/dcl1_1783nt-dsRNA provoked a 79% reduction of *FgDCL1* transcripts. Target gene silencing for *FgDCL2* was also highly efficient, as use of all three constructs, ago1/dcl2_1528nt, ago2/dcl2_1741nt and dcl1/dcl2_1782nt, resulted in an approximately 60% silencing efficiency (Figure 2B). The most efficient construct in terms of overall target gene silencing was dcl1/dcl2_1782nt, which reduced the transcripts of *FgDCL1* and *FgDCL2* by 78% and 58%, respectively, compared to control. Overall, these results suggest that silencing conferred by AGO- and DCL-dsRNAs exhibited the highest efficiency for silencing of *FgDCL1* (AVE: 70%), followed by *FgDCL2* (AVE: 58%), *FgAGO2* (AVE: 48%) and *FgAGO1* (AVE: 26%) (Table 1).

**Table 1:** Overview of target gene-silencing efficiencies of different tested AGO- and DCL-dsRNA constructs.

|  |  | *FgAGO1* | *FgAGO2* | *FgDCL1* | *FgDCL2* |
| --- | --- | --- | --- | --- | --- |
| Tool | *AGO1-DCL1* | 6 | - | 47 | - |
|  | *AGO1-DCL2* | no silencing | - | - | 44 |
|  | *AGO2-DCL1* | - | 41 | 73 | - |
|  | *AGO1-AGO2* | no silencing | 52 | - | - |
|  | **Average** | **6** | **46** | **60** | **44** |
| Manual | *AGO1-DCL1* | 17 | - | 74 | - |
|  | *AGO1-DCL2* | 29 | - | - | 64 |
|  | *AGO2-DCL1* | - | 35 | 79 | - |
|  | *AGO2-DCL2* | - | 49 | - | 67 |
|  | *AGO1-AGO2* | 50 | 62 | - | - |
|  | *DCL1-DCL2* | - | - | 78 | 58 |
|  | **Average** | **32** | **49** | **77** | **63** |

### Manually-designed dsRNAs exhibit higher gene-silencing efficiencies than tool-designed dsRNAs

To assess whether tool-designed dsRNA is more efficient than manually designed constructs, we directly compared target gene-silencing efficiencies of both design approaches (Figure 3). We observed that target gene silencing of manually-designed constructs was superior to tool-designed dsRNA (Figure 3)—except for *FgAGO2*, for which we found no differences between tool- or manually-designed dsRNA. Based on these findings and previous results, we anticipate that larger dsRNA constructs resulted in higher numbers of efficient siRNAs (Höfle et al., 2019a,b). As the tool-designed constructs were <200 nt in length compared to >650 nt for the manually-designed dsRNA (Table 2; Figure S1-4), we calculated possible siRNA hits in the *FgAGO and FgDCL* target genes for all tested dsRNA constructs (Table 2).

**Table 2:**
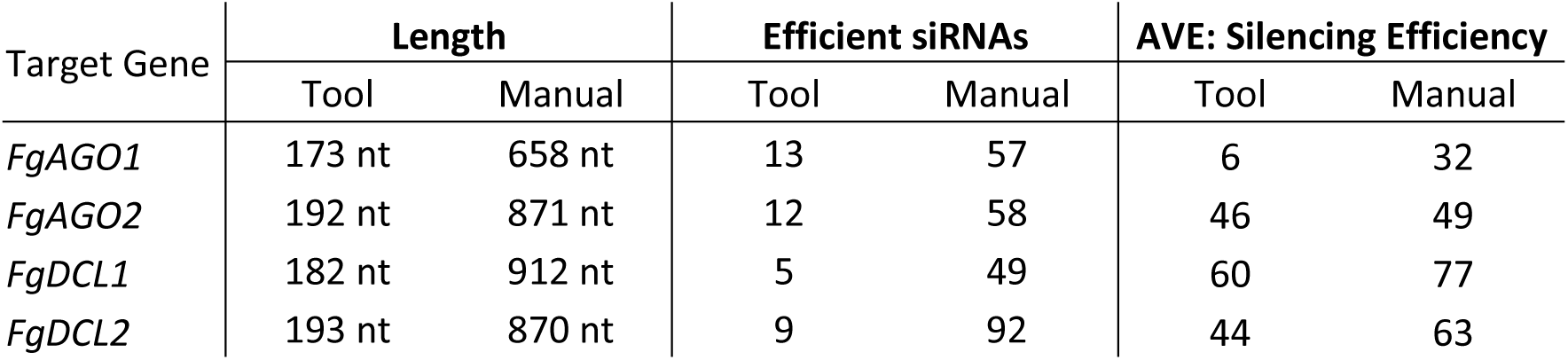
Indicator of how many sRNAs from each double construct fulfil the requirements for efficient target silencing in target genes. These efficient sRNA are designated by the dsRNA design tool si-Fi (http://labtools.ipk-gatersleben.de).

**Figure 3:**
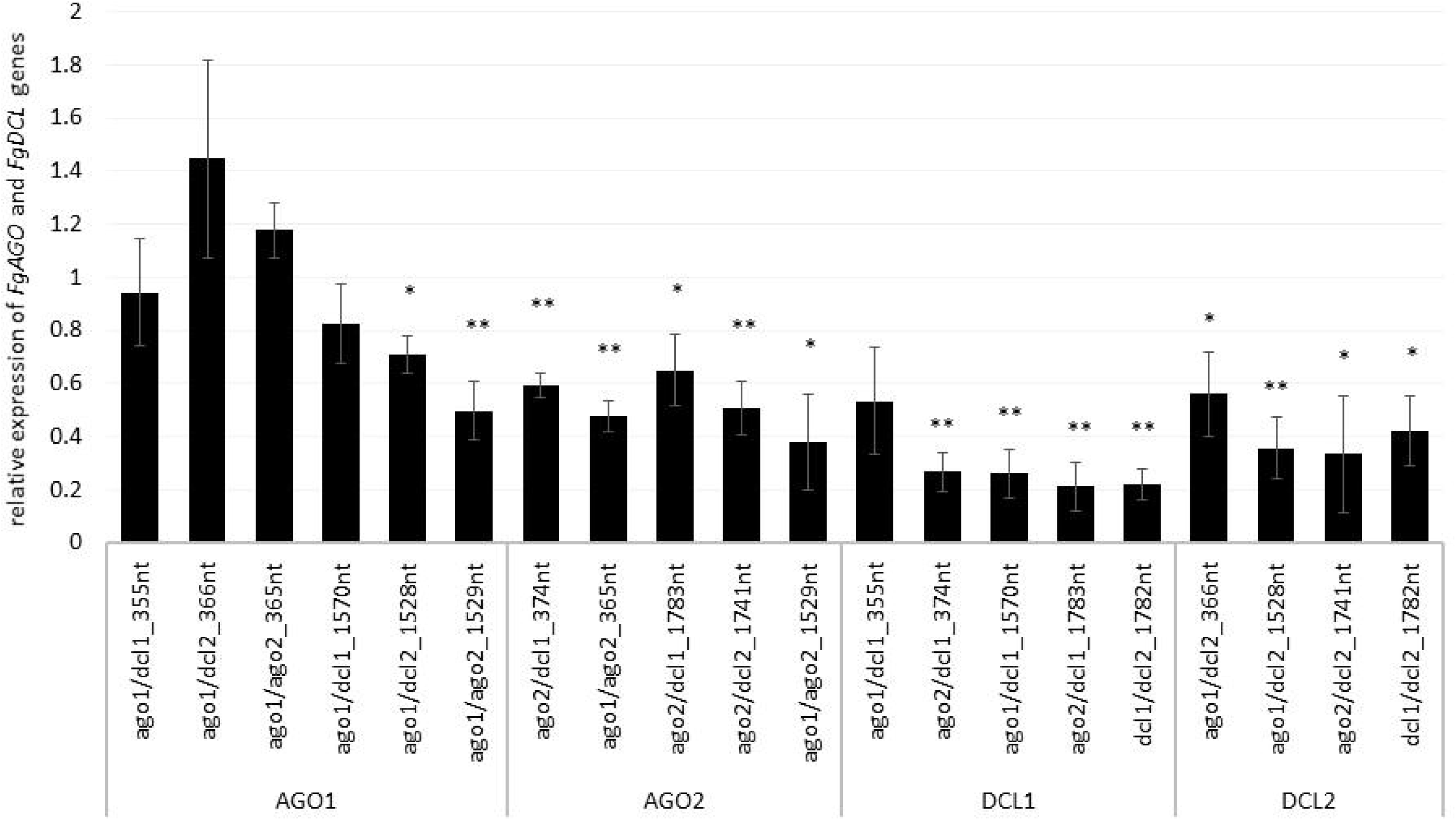
Direct comparison of long (manual) and short (tool) constructs. Relative expression of the respective fungal *DCLs* and *AGOs* 5 dpi on dsRNA-sprayed leaves is grouped by the target gene. The expression was measured via the ΔΔ-ct method in which the expression of the respective *AGOs* and *DCLs* was normalized against the fungal reference gene *EF1α (translation elongation-factor 1 α)* and *ß-tubulin*, and this Δ-ct value was then normalized against the Δ-ct of the GFP control. The asterisks indicate a significant expression of the sprayed leaves in comparison to the mock-treated TE controls. Bars represent mean values ± SE of the four independent experiments.

For the manually-designed dsRNA, we calculated siRNAs that were 4- to 10-fold more efficient compared to the tool-designed constructs (Table 2), thus underlining that the dsRNA precursor length probably plays a role in determining the number of derived siRNAs. For example, we predicted 49 efficient siRNAs were derived out of the 912-nt manually-designed dsRNA, which targets FgDCL1, which is 10-fold greater than the 5 siRNA hits derived from the 182-nt tool-designed *FgDCL1*-dsRNA (Table 2). Notably, these differences resulted in only an overall 10% silencing efficiency decrease of the tool-designed dsRNA compared to the manually-designed constructs targeting *FgDCL1* (Table 2). Together, these data suggest that longer dsRNAs result in a higher number of efficient siRNAs, but there is no stringent correlation that reflects the 10-fold higher number of siRNA resulting in a 10-fold increase in target gene silencing (Table 2).

## Discussion

Microbial pathogens and pests, unlike mammals, are amenable to environmental sRNAs, meaning that they can take up noncoding RNAs from the environment, and these RNAs maintain their RNAi activity (Winston et al. 2007; Whangbo and Hunter 2008; McEwan et al. 2012). This knowledge raises the possibility that plants can be protected from pathogens/pests by exogenously supplied RNA biopesticides (for review, see Mitter et al. 2017; Cai et al. 2018; Dubrovina and Kiselev 2019; Gaffar and Koch 2019; Dalakouras et al. 2020). Possible agronomic application of environmental RNA is affirmed by the high sensitivity of *Fg* to dsRNAs and siRNAs (Koch et al., 2016). Here, we demonstrated that targeting key components of the *Fg* RNAi machinery, such as *AGO* and *DCL* genes, via SIGS could protect barley leaves from *Fg* infection. Our findings, together with other reports, underline that *Fg* RNAi pathways play a crucial role in regulating fungal development, growth, reproduction, mycotoxin production and pathogenicity (Kim et al. 2015; Son et al. 2017; Gaffar et al. 2019). However, the mechanistic roles of *Fg* RNAi components in these processes are inadequately understood. Nevertheless, existing data suggest that there is a functional diversification of *Fg*AGO1/*Fg*DCL2- and *Fg*AGO2/*Fg*DCL1-regulated pathways (Chen et al. 2015; Son et al. 2017; Zeng et al. 2018; Gaffar et al. 2019).

Based on these findings, the dsRNAs tested in this study were designed to target *FgAGO* and *FgDCL* genes pairwise. Thus, we generated six different dsRNA constructs covering all possible *AGO-DCL*-combinations (Fig. 4). Spraying the different dsRNAs onto barley leaves resulted in approximately 50% inhibition of fungal infection for all constructs (Fig. 1). By analysing the silencing efficiencies of the different dsRNA constructs, we found that the expression of *FgDCLs* genes was more suppressed than *FgAGOs* genes (Tab. 1). More importantly, the expression of *FgAGO1* was completely unaffected, regardless of which dsRNA was sprayed. Based on this result, we speculate that *Fg*AGO1 is required for binding of SIGS-associated sRNAs; thus, loss of function mediated by SIGS will not work. Of note, *Δago1* mutants of *Fg* were only slightly compromised in SIGS and less sensitive to dsRNA treatments, indicating redundant functions of *Fg*AGO1 and *Fg*AGO2 in the binding of SIGS-derived siRNAs (Gaffar et al. 2019). However, further studies must explore the mechanistic role of *Fg*AGO1 in SIGS.

**Figure 4:**
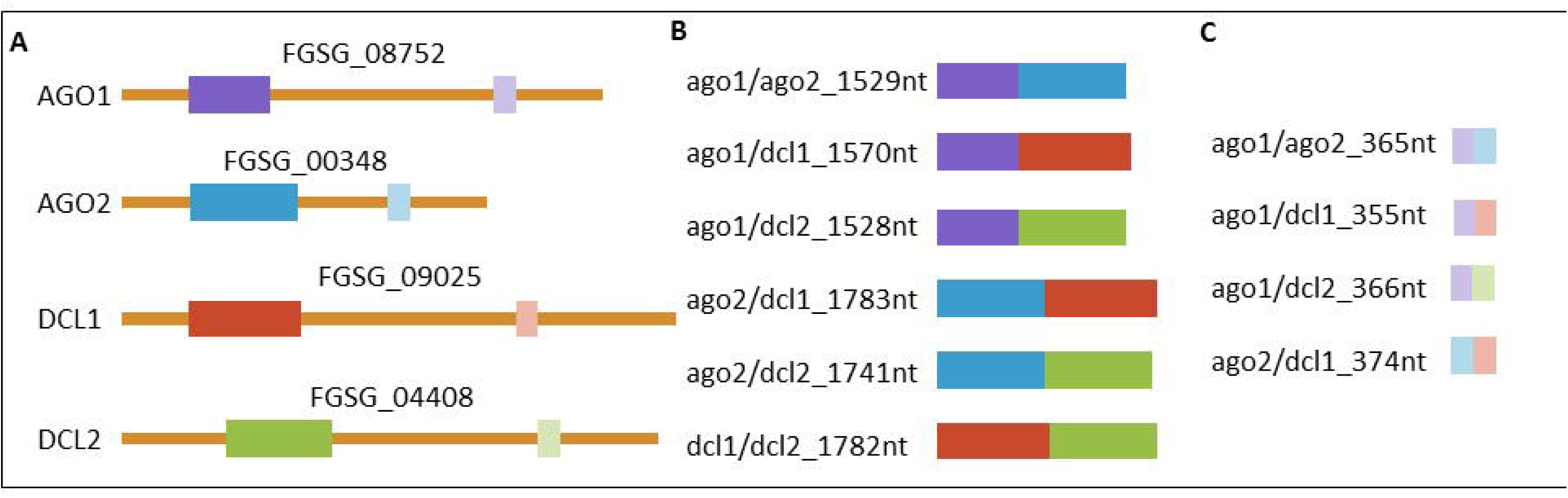
Representation of dsRNAs and the complementary region in the corresponding genes. (A) Graphic representation of all four targeted mRNAs and their respective accessions with target regions marked in colours. Manually selected regions are marked in dark colours and regions selected by the pssRNAit tool (http://plantgrn.noble.org/pssRNAit/) for better target accessibility are marked with light colours. All (B) manually and (C) tool designed dsRNAs triggers are shown. RNAs are represented correctly scaled to each other.

While our data showed that SIGS-mediated downregulation of *FgDCLs* gene expression resulted in inhibition of *Fg* infection, we additionally assume—based on this result—the uptake of siRNAs that were produced by plant DCLs-mediated processing of sprayed dsRNA. Consistent with this finding, previous studies demonstrated that spraying of siRNAs led to the induction of local and systemic RNAi in plants (e.g. Dalakouras et al. 2016; Koch et al. 2016). These findings are significant contributions to our mechanistic understanding of RNA spray technology, as our previous data indicate that SIGS requires the processing of dsRNAs by the fungal RNAi machinery (Koch et al. 2016; Gaffar et al. 2019). Whereas HIGS relies on the host plant’s ability to produce mobile siRNAs (generated from transgene-derived dsRNAs), the mechanism of gene silencing by exogenously delivered dsRNA constitutes a more complex situation; for instance, the possible involvement of the silencing machinery of the host and/or pathogen (Fig. 5). Our previous finding that unprocessed long dsRNA is absorbed from leaf tissue (Koch et al. 2016) has important implications for future disease control strategies based on dsRNA. It is very likely that the application of longer dsRNAs might be more efficient than the application of siRNAs, given their more efficient translocation (Koch et al. 2016). Moreover, in contrast to using only one specific siRNA, processing of long dsRNA into many different inhibitory siRNAs by the fungus may reduce the chance of pathogen resistance under field test conditions. However, RNAi-based plant protection technologies are limited by the uptake of RNAi-inducing trigger molecules, either siRNAs and/or dsRNAs, whereas both RNA types have been shown to confer plant disease resistance independent of how they were applied/delivered (i.e. endogenously or exogenously).

**Figure 5:**
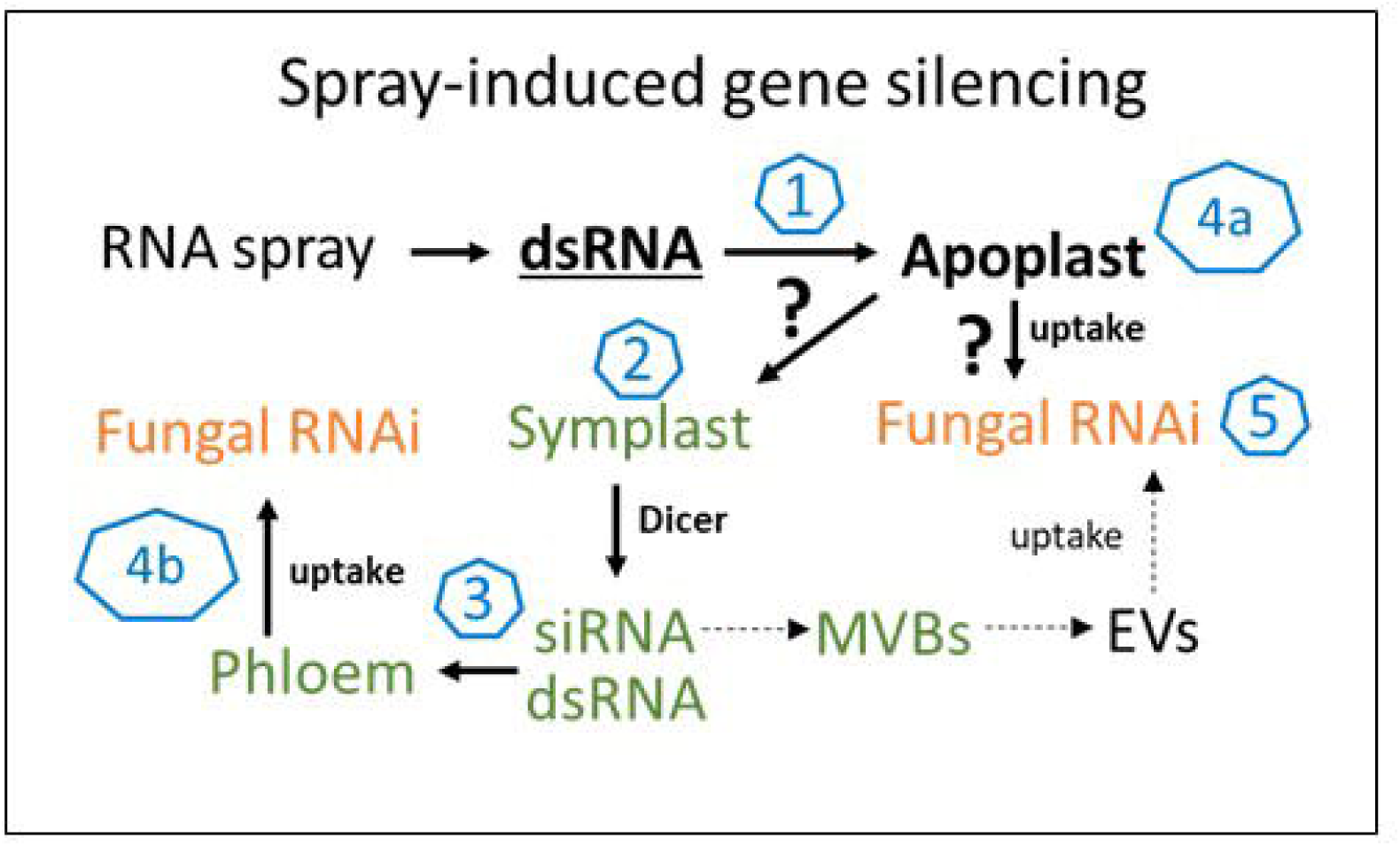
The molecular mechanism of SIGS is controlled by the fungal silencing machinery. In summary, our findings support the model that SIGS involves: (**1**) uptake of sprayed dsRNA by the plant (via stomata); (**2**) transfer of apoplastic dsRNAs into the symplast (DCL processing into siRNAs); (**3**) systemic translocation of siRNA or unprocessed dsRNA via the vascular system (phloem/xylem); (**4**) uptake of apoplastic dsRNA (**a**) or symplastic dsRNA/siRNA by the fungus (**b**); (**5**) processing into siRNA by fungal DCL.

Previously, we discovered that longer dsRNAs of 400–800 nt exhibited a higher gene-silencing efficiency and a stronger disease resistance than 200-nt dsRNAs (Koch et al., 2019), indicating that the quantity of siRNAs derived from a longer dsRNA precursor is simply higher. To test whether the length and/or the selected target gene sequence influences silencing efficiencies, we constructed 10 different dsRNA constructs targeting *FgAGO/FgDCL* pairs (Fig. 4). For the design of the dsRNA constructs we used a dsRNA design tool (http://plantgrn.noble.org/pssRNAit/) that generates dsRNAs of shorter lengths (173-197 nt), compared them to manually selected sequences (658-912 nt) and calculated the number of efficient siRNAs for each construct using si-Fi 2.1 (http://labtools.ipk-gatersleben.de/) *in silico* prediction tool (Table 2). Notably, we found that the number of efficient siRNAs derived from the longer, manually-designed dsRNAs was 4- to 5-fold higher for the constructs that target *FgAGO1* and *FgAGO2*. Moreover, the manually-designed constructs targeting *FgDCL1* and *FgDCL2* resulted in 10-fold more efficient siRNAs than the tool-designed versions (Table 2). However, such a correlation was only observed when we compared tool-versus manually-designed dsRNA—<200-nt versus >650-nt constructs. If we attempt to predict the number of efficient siRNAs of all the manually designed dsRNAs based on the length of their precursors, we obtain contrasting results. For example, the 912-nt precursor dsRNA that targets *FgDCL1* resulted in 49 efficient siRNA hits, which is approximately half of the 92 siRNA hits for the 870-nt dsRNA designed to target *FgDCL2* (Table 2). Importantly, the tested dsRNAs that target *FgDCL1*, which showed the lowest number of siRNAs, revealed the highest efficiencies compared to all other constructs (Table 2). Together, our data support the notion that longer dsRNAs tend to result in higher numbers of siRNA, although this can differ in particular cases.

However, these data were obtained from *in silico* predictions; therefore, their accuracies remain unknown. As such, RNA-sequencing must be performed to quantify, analyse and map the SIGS-derived siRNAs to their target genes as well as their dsRNA precursors. Nevertheless, besides their concentration, it is known that the siRNA sequence represents a crucial determinant regarding silencing efficiencies of their complementary target genes (Ossowski et al. 2008). In addition, mapping siRNAs to their target sequence revealed processing patterns that might help to define principles for RNAi vector design, producing effective siRNAs (Yang et al. 2013; Koch et al. 2016; Baldwin et al. 2018). Importantly, to construct our manually-designed dsRNAs, we performed a random selection of sequences complementary to the specific target genes. Moreover, to guarantee optimal silencing, we chose longer dsRNA sequences compared to the tool-designed dsRNAs which along with targeting two genes (*AGO/DCL* pairs), resulted in relatively long dsRNAs (>1kb). Thus, a random selection of longer target sequences tends to increase off-target effects per se (Roberts et al. 2015). Therefore, based on our results obtained with tool-designed dsRNAs and the work of others, we suggest using minimal-length dsRNA sequences carefully selected based on identified determining design criteria. Another possible way to achieve high silencing efficiencies while retaining high target specificity could be the construction of dsRNAs repeating a shorter tool-designed sequence several times.

Nevertheless, the number of efficient siRNAs that reach the fungus depends on the uptake efficiency of sprayed dsRNA molecules and that can differ depending on the parameters which determine an efficient uptake, such as the stomata opening (Koch et al. 2016). Additionally, the concentration of siRNAs in the target organisms can vary, as we previously found that SIGS mainly rely on the uptake of unprocessed dsRNA from the plant’s apoplast and their processing by fungal DCLs (Koch et al. 2016; Gaffar et al. 2019). Finally, and even more important than quantities of target-specific siRNAs in determining silencing efficacy, is the target accessibility of a siRNA (Reynolds et al. 2004; Shao et al. 2007). Therefore, the design of RNAi constructs that likely mediate the efficient uptake of dsRNAs and/or siRNAs by the target pathogen is crucial in determining the success of SIGS as well as HIGS technologies.

Together, our results indicate that silencing fungal RNAi pathway genes, especially *DCL* genes, using SIGS efficiently increases plant disease resistance towards necrotrophic fungal pathogens, such as *Fg*. Moreover, our results support the notion that fungal RNAi-related genes in *Fg* play an essential role in pathogenicity and/or virulence (Gaffar et al., 2019). These findings are consistent with other reports demonstrating that the two DCL proteins (DCL1 and DCL2) of the fungal pathogen *Botrytis cinerea* (*Bc*) play a central role in disease development (Wang et al. 2016). The authors show that the application of sRNAs or dsRNAs on fruits, vegetables and flowers targeting *BcDCL1* and *BcDCL2* genes significantly inhibited grey mould disease. Of note, the same group previously discovered that *Bc* delivers sRNAs into plant cells to silence host immunity genes, a phenomenon called ‘cross-kingdom RNAi (ckRNAi)’ (Weiberg et al. 2013). Emerging data further suggest that some sRNA effectors can target multiple host defence genes to enhance *Bc* pathogenicity. For example, Bc-siR37 suppresses host immunity by targeting at least 15 *Arabidopsis* genes, including WRKY transcription factors, receptor-like kinases and cell wall-modifying enzymes (Wang et al., 2017a). Moreover, one of the most destructive pathogens of wheat *Puccinia striiformis* also delivers fungal sRNAs, such as microRNA-like RNA1 (milR1), into host cells and suppresses wheat *Pathogenesis-related 2* (*PR-2*) in the defence pathway (Wang et al., 2017b). Notably, such ckRNAi-related sRNA effectors are produced by DCL proteins, and thus SIGS of fungal DCLs abolishes sRNA production and attenuates fungal pathogenicity and growth. However, whether our findings suggest that *Fg* utilizes ckRNAi-related sRNAs to suppress host immunity needs further exploration.

More importantly, while several studies have demonstrated bidirectional ckRNAi and sRNA trafficking between plant hosts and their interacting fungal pathogens (Zhang et al., 2012; Weiberg et al., 2013; Zhang et al., 2016; Wang et al., 2017a/b; Zhu et al., 2017; Cai et al., 2018; Dubey et al., 2019; Zanini et al., 2019), the mechanisms underlying the transfer and uptake of transgene-derived artificial sRNAs (HIGS) as well as exogenously applied dsRNA (SIGS) remain elusive. Further research is needed to determine, for example: (i) how plant and fungal-silencing machinery contributes to HIGS and SIGS; (ii) the nature of the inhibitory RNA that translocates from the plant to the fungus after its transgenic expression or spray application; (iii) how that RNA crosses the plant-fungal interface; and (iv) how dsRNA is transported at the apoplast-symplast interface. Therefore, addressing these questions is key for making RNA silencing-based strategies a realistic and sustainable approach in agriculture.

## Acknowledgments

This work was supported by Deutsche Forschungsgemeinschaft to AK (RTG:2355) and Deutscher Akademischer Austauschdienst to FYG.

## Figure legends

**Supplemental Figure 1-4**: Coding sequences (CDS) of the respective *Fg* target gene with the sequences of the dsRNA marked (blue: tool-designed; red: manually designed).

**Supplemental Table 1:** Primer used in this study

**Supplemental Figure 2:** Representative pictures of barley (golden promise) leaves sprayed with 10µg (20ng/µl) of respective dsRNA in TE-Buffer and the control w/o dsRNA. dsRNA was applied on the upper half of 10 leaves and 2 days after spraying the leaves were inoculated with three 20µl droplets of *Fg* (50.000 spores/ml). The pictures were taken 5dpi.

## References

Abdellatef, E., Will, T., Koch, A., Imani, J., Vilcinskas, A. and Kogel, K.H. (2015) Silencing the expression of the salivary sheath protein causes transgenerational feeding suppression in the aphid *Sitobion avenae*. Plant Biotechnology Journal, 849–57. doi: 10.1111/pbi.12322.

Baldwin, T., Islamovic, E., Klos, K., Schwartz, P., Gillespie, J., Hunter, S., & Bregitzer, P. (2018). Silencing efficiency of dsRNA fragments targeting Fusarium graminearum TRI6 and patterns of small interfering RNA associated with reduced virulence and mycotoxin production. PloS one, 13(8), e0202798.

Baulcombe, D. (2004). RNA silencing in plants. Nature, 431(7006), 356.

Bharti, P., Jyoti, P., Kapoor, P., Sharma, V., Shanmugam, V., & Yadav, S. K. (2017). Host-induced silencing of pathogenicity genes enhances resistance to Fusarium oxysporum wilt in tomato. Molecular biotechnology, 59(8), 343–352.

Borges, F., & Martienssen, R. A. (2015). The expanding world of small RNAs in plants. Nature reviews Molecular cell biology, 16(12), 727.

Cai, Q., Qiaom L., Wang, M., He, B., Lin, FM., Palmquist, J., Huang, SD. and Jin, H. (2018) Plants send small RNAs in extracellular vesicles to fungal pathogen to silence virulence genes. Science. 1126–1129. doi: 10.1126/science.aar4142.

Cai, Q., He, B., Kogel, K.H., and Jin, H. (2018). Cross-kingdom RNA trafficking and environmental RNAi nature’s blueprint for modern crop protection strategies. Current Opinion Microbiology, 46, 58–64. doi: 10.1016/j.mib.2018.02.003.

Castel, S.E. and Martienssen R.A. (2013) RNA interference in the nucleus: roles for small RNAs in transcription, epigenetics and beyond. Nat. Rev. Genet. 14,100–112. doi: 10.1038/nrg3355.

Chen, Y., Gao, Q., Huang, M., Liu, Y., Liu, Z., Liu, X., & Ma, Z. (2015). Characterization of RNA silencing components in the plant pathogenic fungus Fusarium graminearum. Scientific reports, 5, 12500.

Chen, W., Kastner, C., Nowara, D., Oliveira-Garcia, E., Rutten, T., Zhao, Y., … & Schweizer, P. (2016). Host-induced silencing of Fusarium culmorum genes protects wheat from infection. Journal of experimental botany, 67(17), 4979–4991.

Cheng, W., Song, X. S., Li, H. P., Cao, L. H., Sun, K., Qiu, X. L., … & Qu, B. (2015). Host-induced gene silencing of an essential chitin synthase gene confers durable resistance to F usarium head blight and seedling blight in wheat. Plant biotechnology journal, 13(9), 1335–1345.

Dalakouras, A., Wassenegger, M., Dadami, E., Ganopoulos, I., Pappas, M. L., & Papadopoulou, K. Genetically Modiied Organism-Free RNA Interference: Exogenous. Plant Physiol, 182, 2020.

Dalakouras, A., Wassenegger, M., McMillan, J. N., Cardoza, V., Maegele, I., Dadami, E., … & Wassenegger, M. (2016). Induction of silencing in plants by high-pressure spraying of in vitro-synthesized small RNAs. Frontiers in Plant Science, 7, 1327.

Dubey, H., Kiran, K., Jaswal, R., Jain, P., Kayastha, A.M., Bhardwaj, S.C., Mondal, T.K. and Sharma T.R. Discovery and profiling of small RNAs from Puccinia triticina by deep sequencing and identification of their potential targets in wheat. Funct Integr Genomics. 391–407. doi: 10.1007/s10142-018-00652-1.

Dubrovina, A. S., & Kiselev, K. V. (2019). Exogenous RNAs for Gene Regulation and Plant Resistance. International journal of molecular sciences, 20(9), 2282.

Gaffar, F.Y. and Koch, A. (2019) Catch me if you can! RNA silencing-based improvement of antiviral plant immunity. Under revision.

Gaffar, F.Y., Imani, J., Karlovsky, J.P., Koch, A. and Kogel, K.H. (2019) Various components of the RNAi pathway are required for conidiation, ascosporogenesis, virulence, DON production and SIGS-mediated fungal inhibition by exogenous dsRNA in the Head Blight pathogen *Fusarium graminearum*. Frontiers in Microbiology, in press.

Ghag, S. B., Shekhawat, U. K., & Ganapathi, T. R. (2014). Host-induced post-transcriptional hairpin RNA-mediated gene silencing of vital fungal genes confers efficient resistance against F usarium wilt in banana. Plant biotechnology journal, 12(5), 541–553.

Goswami, R. S., & Kistler, H. C. (2004). Heading for disaster: Fusarium graminearum on cereal crops. Molecular plant pathology, 5(6), 515–525.

Guo, Q., Liu, Q., Smith, N.A., Liang, G., and Wang, M.B. (2016) RNA Silencing in plants: mechanisms, technologies and applications in horticultural crops. Current Genomics, 17, 476–489. doi: 10.2174/1389202917666160520103117.

Höfle, L., Koch, A., Schmidt, A., Imani, J., Jelonek, L. and Kogel, K.H. (2019) SIGS vs HIGS: A comparative study on the efficacy of antifungal double-stranded RNAs targeting *Fusarium FgCYP51* genes. Under review.

Höfle, L., Shrestha, A., Jelonek, L. and Koch, A. (2019) Study on the efficacy of dsRNAs with increasing length targeting *Fusarium graminearum CYP51* genes comparing HIGS and SIGS approaches. Under revision.

Hu, Z., Parekh, U., Maruta, N., Trusov, Y., & Botella, J. R. (2015). Down-regulation of Fusarium oxysporum endogenous genes by host-delivered RNA interference enhances disease resistance. Frontiers in chemistry, 3, 1.

Ismaiel, A., & Papenbrock, J. (2015). Mycotoxins: producing fungi and mechanisms of phytotoxicity. Agriculture, 5(3), 492–537.

Kaldis, A., Berbati, M., Melita, O., Reppa, C., Holeva, M., Otten, P. and Voloudakis, A. (2018) Exogenously applied dsRNA molecules deriving from the Zucchini yellow mosaic virus (ZYMV) genome move systemically and protect cucurbits against ZYMV. Mol Plant Pathol. 883–895. doi: 10.1111/mpp.12572.

Kazan, K., Gardiner, D. M., & Manners, J. M. (2012). On the trail of a cereal killer: recent advances in Fusarium graminearum pathogenomics and host resistance. Molecular plant pathology, 13(4), 399–413.

Ketting, R. F. (2011). The many faces of RNAi. Developmental cell, 20(2), 148–161.

Kim, H. K., Jo, S. M., Kim, G. Y., Kim, D. W., Kim, Y. K., & Yun, S. H. (2015). A large-scale functional analysis of putative target genes of mating-type loci provides insight into the regulation of sexual development of the cereal pathogen Fusarium graminearum. PLoS genetics, 11(9), e1005486.

Koch, A., Kang, H.G., Steinbrenner, J., Dempsey, D.A., Klessig, D.F. and Kogel, K.H. (2017). MORC Proteins: Novel Players in Plant and Animal Health. Frontiers in Plant Science. doi.org/10.3389/fpls.2017.01720

Koch, A. and Kogel, K.-H. (2014). New wind in the sails: improving the agronomic value of crop plants through RNAi-mediated gene silencing. Plant Biotechnology Journal, 821–831, doi: 10.1111/pbi.12226

Koch, A., Biedenkopf, B., Furch, A.C.U., Abdellatef, E., Weber, L., Linicus, L., Johannsmeier, J., Jelonek, L., Goesmann, A., Cardoza, V., McMillan, J., Mentzel, T. and Kogel, K.H. (2016). An RNAi-based control of Fusarium graminearum infections through spraying of long dsRNAs involves a plant passage and is controlled by the fungal silencing machinery Plos Pathogens, 12, doi: 10.1371/journal.ppat.1005901.

Koch, A., Kumar, N., Weber, L., Keller, H., Imani, J. and Kogel, K.H. (2013). Host-induced gene silencing of cytochrome P450 lanosterol C14α-demethylase-encoding genes confers strong resistance to Fusarium spec. PNAS, doi: 10.1073/pnas.1306373110.

Koch, A., Stein, E., & Kogel, K. H. (2018). RNA-based disease control as a complementary measure to fight Fusarium fungi through silencing of the azole target Cytochrome P450 Lanosterol C-14 α-Demethylase. European journal of plant pathology, 152(4), 1003–1010.

Koch A, Höfle L, Werner BT, Imani J, Schmidt A, Jelonek L, Kogel K. H. (2019). SIGS vs HIGS: A study on the efficacy of two dsRNA delivery strategies to silence Fusarium FgCYP51 genes in infected host and non-host plants. Molecular Plant Pathology, in press

Konakalla, N.C., Kaldis, A., Berbati, M., Masarapu, H. and Voloudakis, A.E. (2016). Exogenous application of double-stranded RNA molecules from TMV p126 and CP genes confers resistance against TMV in tobacco. Planta. 961–9. doi: 10.1007/s00425-016-2567-6.

Kuck, K. H., Stenzel, K., & Vors, J. P. (2012). Sterol biosynthesis inhibitors. Modern crop protection compounds, 1, 761–805.

Livak, K. J., & Schmittgen, T. D. (2001). Analysis of relative gene expression data using real-time quantitative PCR and the 2− ΔΔCT method. methods, 25(4), 402–408.

Machado, A. K., Brown, N. A., Urban, M., Kanyuka, K., & Hammond-Kosack, K. E. (2018). RNAi as an emerging approach to control Fusarium head blight disease and mycotoxin contamination in cereals. Pest management science, 74(4), 790–799.

Majumdar, R., Rajasekaran, K., & Cary, J. W. (2017). RNA interference (RNAi) as a potential tool for control of mycotoxin contamination in crop plants: concepts and considerations. Frontiers in plant science, 8, 200.

McEwan, D. L., Weisman, A. S., & Hunter, C. P. (2012). Uptake of extracellular double-stranded RNA by SID-2. Molecular cell, 47(5), 746–754.

McMullen, M., Bergstrom, G., De Wolf, E., Dill-Macky, R., Hershman, D., Shaner, G., & Van Sanford, D. (2012). A unified effort to fight an enemy of wheat and barley: Fusarium head blight. Plant Disease, 96(12), 1712–1728.

Mitter, N., Worrall, E.A., Robinson, K.E., Li, P., Jain, R.G., Taochy, C., Fletcher, S.J., Carroll, B.J., Lu, G.Q. and Xu, Z.P. (2017). Clay nanosheets for topical delivery of RNAi for sustained protection against plant viruses. Nature plants 3, 16207. doi: 10.1038/nplants.2016.207.

Nowara, D., Gay, A., Lacomme, C., Shaw, J., Ridout, C., Douchkov, D., Hensel, G., Kumlehn J., and Schweizer, P. (2010). HIGS: host-induced gene silencing in the obligate biotrophic fungal pathogen Blumeria graminis. The Plant Cell 3130–3141, oi: 10.1105/tpc.110.077040.

Pareek, M., & Rajam, M. V. (2017). RNAi-mediated silencing of MAP kinase signalling genes (Fmk1, Hog1, and Pbs2) in Fusarium oxysporum reduces pathogenesis on tomato plants. Fungal biology, 121(9), 775–784.

Qi, T., Guo, J., Peng, H., Liu, P., Kang, Z. and Guo, J. (2019). Host-Induced Gene Silencing: A Powerful Strategy to Control Diseases of Wheat and Barley. Int J Mol Sci. doi: 10.3390/ijms20010206.

Reynolds, A., Leake, D., Boese, Q., Scaringe, S., Marshall, W. S., & Khvorova, A. (2004). Rational siRNA design for RNA interference. Nature biotechnology, 22(3), 326.

Roberts, A. F., Devos, Y., Lemgo, G. N., & Zhou, X. (2015). Biosafety research for non-target organism risk assessment of RNAi-based GE plants. Frontiers in plant science, 6, 958.

Schneider, C. A., Rasband, W. S., & Eliceiri, K. W. (2012). NIH Image to ImageJ: 25 years of image analysis. Nature methods, 9(7), 671.

Shao, Y., Chan, C. Y., Maliyekkel, A., Lawrence, C. E., Roninson, I. B., & Ding, Y. (2007). Effect of target secondary structure on RNAi efficiency. Rna, 13(10), 1631–1640.

Son, H., Park, A. R., Lim, J. Y., Shin, C., & Lee, Y. W. (2017). Genome-wide exonic small interference RNA-mediated gene silencing regulates sexual reproduction in the homothallic fungus Fusarium graminearum. PLoS genetics, 13(2), e1006595.

Spolti, P., Del Ponte, E. M., Dong, Y., Cummings, J. A., & Bergstrom, G. C. (2014). Triazole sensitivity in a contemporary population of Fusarium graminearum from New York wheat and competitiveness of a tebuconazole-resistant isolate. Plant disease, 98(5), 607–613.

Vaucheret, H., Vazquez, F., Crété, P., & Bartel, D. P. (2004). The action of ARGONAUTE1 in the miRNA pathway and its regulation by the miRNA pathway are crucial for plant development. Genes & development, 18(10), 1187–1197.

Wang, B., Sun, Y.F., Song, N., Zhao, M.X., Liu, R., Feng, H., Wang, X.J. and Kang, Z.S. (2017b). *Puccinia striiformis* f. sp *tritici* microRNA-like RNA 1 (Pst-milR1), an important pathogenicity factor of Pst, impairs wheat resistance to Pst by suppressing the wheat pathogenesis-related 2 gene. New Phytologist, 338–350. doi: 10.1111/nph.14577.

Wang, M., Weiberg, A., Dellota, E. Jr, Yamane, D, and Jin, H. (2017a). Botrytis small RNA Bc-siR37 suppresses plant defense genes by cross-kingdom RNAi. RNA Biology, 421–428. doi: 10.1080/15476286.2017.

Wang, M., Weiberg, A., Lin, F.-M., Thomma, B. P. H. J., Huang, H.D., and Jin, H. (2016). Bidirectional cross-kingdom RNAi and fungal uptake of external RNAs confer plant protection. Nature Plants, 2, 16151. doi: 10.1038/nplants.2016.151.

Weiberg, A., Bellinger, M. and Jin, H.L. (2015). Conversations between kingdoms: small RNAs. Current Opinion Biotechnology, 207–215. doi: 10.1016/j.copbio.2014.

Weiberg, A., Wang, M., Lin, F.M., Zhao, H., Zhang, Z., Kaloshian, I., Huang, H.D. and Jin, H. (2013). Fungal small RNAs suppress plant immunity by hijacking host RNA interference pathways. Science, 118–123. doi: 10.1126/science.1239705.

Whangbo, J. S., & Hunter, C. P. (2008). Environmental RNA interference. Trends in genetics, 24(6), 297–305.

Winston, W. M., Sutherlin, M., Wright, A. J., Feinberg, E. H., & Hunter, C. P. (2007). Caenorhabditis elegans SID-2 is required for environmental RNA interference. Proceedings of the National Academy of Sciences, 104(25), 10565–10570.

Yin, C. and Hulbert, S. (2015). Host Induced Gene Silencing (HIGS), a Promising Strategy for Developing Disease Resistant Crops. Gene Technology 4:130. doi: 10.4172/2329-6682.1000130

Yin, Y., Liu, X., Li, B., & Ma, Z. (2009). Characterization of sterol demethylation inhibitor-resistant isolates of Fusarium asiaticum and F. graminearum collected from wheat in China. Phytopathology, 99(5), 487–497.

Zanini, S., Šečić, E., Busche, T., Kalinowski, J., & Kogel, K. H. (2019). Discovery of interaction-related sRNAs and their targets in the Brachypodium distachyon and Magnaporthe oryzae pathosystem. BioRxiv, 631945.

Zeng, W., Wang, J., Wang, Y., Lin, J., Fu, Y., Xie, J., … & Cheng, J. (2018). Dicer-like proteins regulate sexual development via the biogenesis of perithecium-specific microRNAs in a plant pathogenic fungus Fusarium graminearum. Frontiers in microbiology, 9, 818.

Zhang, J., Khan, S.A., Heckel, D.G. and Bock, R. (2017). Next-Generation Insect-Resistant Plants: RNAi-Mediated Crop Protection. Trends in biotechnology. doi: 10.1016/j.tibtech.2017.04.009.

Zhang, L., Hu, D., Chen, X., Li, D., Zhu, L., Zhang, Y., Li, J., Bian, Z., Liang, X., Cai, X., Yin, Y., Wang, C., Zhang, T., Zhu, D., Zhang, D., Xu, J., Chen, Q., Ba, Y., Liu, J., Wang, Q., Chen, J., Wang, J., Wang, M., Zhang, Q., Zhang, J., Zen, K. and Zhang, C.Y. (2012). Exogenous plant MIR168a specifically targets mammalian LDLRAP1: evidence of cross-kingdom regulation by microRNA. Cell Res. 107–26. doi: 10.1038/cr.2011.158.

Zhang, T., Zhao, Y.L., Zhao, J.H., Wang, S., Jin, Y., Chen, Z.Q., Fang, Y.Y., Hua, C.L., Ding, S.W., Guo, H.S. (2016). Cotton plants export microRNAs to inhibit virulence gene expression in a fungal pathogen Nat Plants. 16153. doi: 10.1038/nplants.2016.153.

Zhu, K., Liu, M., Fu, Z., Zhou, Z., Kong, Y., Liang, H., Lin, Z., Luo, J., Zheng, H., Wan, P., Zhang, J., Zen, K., Chen, J., Hu, F., Zhang, C.Y., Ren, J., Chen, X. (2017). Plant microRNAs in larval food regulate honeybee caste development. PLoS Genet. e1006946. doi: 10.1371/journal.pgen.1006946.

Mitter, N., Worrall, E. A., Robinson, K. E., Xu, Z. P., & Carroll, B. J. (2017). Induction of virus resistance by exogenous application of double-stranded RNA. Current opinion in virology, 26, 49–55.

Yu, J., Lee, K. M., Cho, W. K., Park, J. Y., & Kim, K. H. (2018). Differential contribution of RNA interference components in response to distinct Fusarium graminearum virus infections. Journal of virology, 92(9), e01756–17.

Ossowski, S., Schwab, R., & Weigel, D. (2008). Gene silencing in plants using artificial microRNAs and other small RNAs. The Plant Journal, 53(4), 674–690.

Yang, Y., Jittayasothorn, Y., Chronis, D., Wang, X., Cousins, P., & Zhong, G. Y. (2013). Molecular characteristics and efficacy of 16D10 siRNAs in inhibiting root-knot nematode infection in transgenic grape hairy roots. PloS one, 8(7), e69463.

